## Supplementary material for "An affordable solution for investigating zebra finch intracranial electroencephalography (iEEG) signals": https://www.protocols.io/view/an-affordable-solution-for-investigating-zebra-fin-q26g7y2w8gwz/v3?version_warning=no&step=12

dkiq4udw

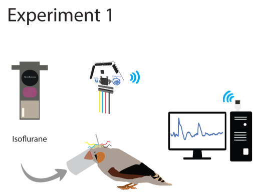


Milad Yekani[1], Mohammad-Mahdi Abolghasemi[2], Shahrar Rezghi Shirsavar[1]

[1]School of Cognitive Sciences, Institute for Research in Fundamental Sciences (IPM), Tehran, Iran;
[2]Institute for Cognitive and Brain Sciences, Shahid Beheshti University, Tehran, Iran.

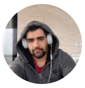

Milad Yekani
IPM cognitive school

**Protocol Info:** Milad Yekani, Mohammad-Mahdi Abolghasemi, Shahrar Rezghi Shirsavar . An affordable solution for investigating zebra finch intracranial EEG signals. **protocols.io** **https://protocols.io/view/an-affordable-solution-for-investigating-zebra-fin-dkiq4udw**

**Created:** May 07, 2023

**Last Modified:** October 28, 2024

**Protocol Integer ID:** 106800

**Keywords:** animal EEG, open source, Song bird, Zebra finch

#### Disclaimer

The SCS Research Ethics Committee of IPM approves all surgical steps presented here. Some of these procedures may not be accepted in other research facilities.

### Abstract

The Zebra finch is a well-studied model animal to study neural mechanisms of speech, and electrophysiology is the primary technique for understanding the song system in them. Most of the studies on zebra finches have focused on intracerebral recordings. However, these methods are only affordable for limited laboratories. Recently, different open-source hardware for acquiring EEG signals has been developed. It's unclear if these solutions are suitable for zebra finch studies as they have not been evaluated so far. Electrocorticography signals can provide a preliminary guide for more in-depth inquiries and also aid in understanding the global behavior of the bird's brain, as opposed to the more common localized approach. We provide a detailed protocol for acquiring iEEG data from zebra finches with an open-source device. We implemented stainless steel electrodes on the brain's surface and recorded

the brain signals from two recording sites. To validate our method, we ran two different experiments. In the first experiment, we recorded neural activity under various concentrations of isoflurane and extracted the suppression duration to measure anesthesia depth. In the second experiment, we head-fixed the birds

and presented them with different auditory stimuli to evaluate event-related potential (ERP). Results showed a significant increase in the suppression duration by increasing the anesthesia depth and evident ERP response to auditory stimuli. These findings indicate that by our methodology, we can successfully collect iEEG signals from awake and anesthetized birds. These findings pave the way for future studies to use iEEG to investigate bird cognition.

### Materials

**########################## in this table we are going to list all materials###################**

"""""""""""""""""""""""""""""""""""""""""""""""" find required materials in your step list in this table""""""""""""""""""""""""""""""""""""

| A | B | C | D |
|---|---|---|---|
| **Material** | **Source** | **Price** | **Number** |
| anesthesia mask | homemeade (3D printed) | 20$ | 1 |
| hypodermic needle | https://www.medirancenter.com/ | 5 $ | 10 |
| stainless steel wire(Catalog: 791500) | https://www.a-msystems.com/p-809-pfa-coated-stainless-steel-wire.aspx | 166$ | 100 cm |
| CNY17F optocoupler | https://www.vishay.com/en/product/83606/ | 0.2$ | 1 |
| Duracell Plus AA Alkaline Batteries | https://www.duracell.co.uk/product/duracell-plus-type-aa-alkaline-batteries/ | ~5$ | 4 |
| Jumper Wires | Amazon | ~5$ | 4 |
| Hook Clip Probe Test | Amazon | ~7$ | 8-10 |
| Raspberry pi 3 | https://www.raspberrypi.com/products/raspberry-pi-3-model-b/ | ~35$ | 1 |
| Bread board | Amazon | ~5$ | 1 |
| Open BCI cyton board | https://shop.openbci.com/products/cyton-biosensing-board-8-channel | ~900$ | 1 |
| 1k ressitor | Amazon | 1$ | 2 |
| C_silicon + activator | coltene | ~ | 1 |
| Isoflurane | | | |
| surgical micromotor | https://www.amazon.in/Oscar-Saeshin-Dental-Polishing-Handpiece/dp/B079W6R1N2 | 90$ | 1 |
| 1 mm ball-shaped dental burrel | | 3$ | 1 |
| Glass inomer cement | http://www.agmdental.ir/sitepages/view-6607.aspx | 15 $ | 1 |
| ciprofloxacin (0.3% eye drop) | Sinadarou Labs Company | 0.5 $ | 1 |

#### Safety warnings

- Isoflurane leakage from anesthesia machines can affect experimenters' health. To avoid this issue, use a suitable scavenging system. Staff should be well informed and educated about the possible hazards ***of anesthetic gas.***
- **¥** marks sections that may require special approval from the local ethical committee.

#### Ethics statement

All procedures were approved by the Research Ethics Committee of the School of Cognitive Sciences (SCS) of the Institute for Research in Fundamental Sciences (IPM, protocol number 1402/40/1/2841).

#### Assembling the recording apparatus

1 **Get started with open BCI:**
Below are essential links for purchasing boards, setup guides, software documentation, and troubleshooting resources for the Cyton and Ganglion boards.
Note: We have used the Cyton board for our project. **However, others can use the Ganglion board as an alternative.**

**Cyton Board Purchase Link:**
Cyton Biosensing Board (8-Channel)
**Ganglion Board (Alternative to Cyton) Purchase Link:**
Ganglion Board
**Getting Started with Cyton Board and OpenBCI GUI:**
Cyton Getting Started Guide
**OpenBCI GUI Guide and Software Download Links:**
OpenBCI GUI Documentation
**Cyton Board External Trigger (Tagging Signals):**
Cyton External Trigger Guide
**Optoisolator Specification and Purchase Links:**
Vishay Optoisolator Specification
Mouser Purchase Link for Vishay Optoisolator
**Troubleshooting (Noise, GUI, and FTDI Buffer Fix):**
OpenBCI Troubleshooting Guide

2 **Testing the hardware:**
To connect the OpenBCI board and start recording, follow these steps:

2.1 **Plug in the OpenBCI USB Dongle:** Insert the dongle, ensuring the blue LED stays on and the red LED blinks. The switch should be set to GPIO 6 (not RESET).

2.2 **Connect the Battery:** Use a Lithium Polymer battery or a 6V AA battery pack. Ensure the voltage is within the specified range to avoid damage.

2.3 **Turn on the Cyton Board:** Switch the board to "PC" mode. The blue LED should light up. If it doesn't, check the battery or press the reset button.

2.4 **Connect the Electrodes:** Attach the electrodes to the bottom pins of the board as follows (you can select any of N1P to N7P).

| A | B |
|---|---|
| **Cyton Board Pin** | **Function** |
| SRB2 (bottom SRB pi n) | Reference Pin |

| A | B |
|---|---|
| bottom BIAS pin | Noise-cancelling Pin |
| 2N (bottom N2P pin) | Analog input |
| 7N (bottom N7P pin) | Analog input |

Table1: Information about the board's pins

3 **Connecting via GUI:**

3.1 Open the GUI and set the data source to LIVE from the Cyton board (Figure 1).

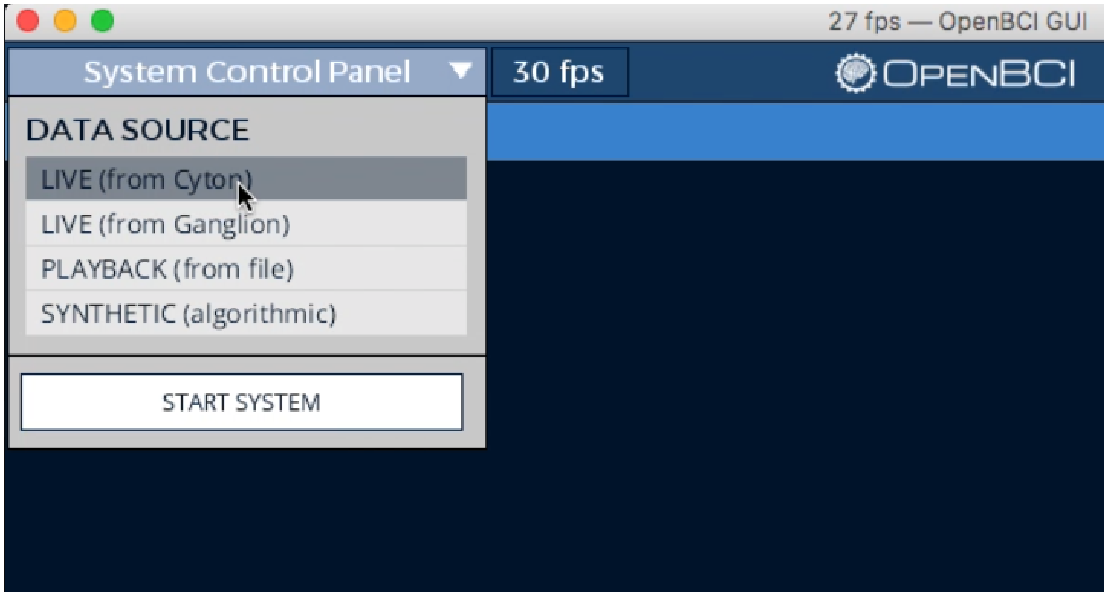


**Figure 1:** In the System Control Panel, click on "LIVE."

3.2 Choose "Serial (from Dongle)" as the data transfer option (Figure 2).

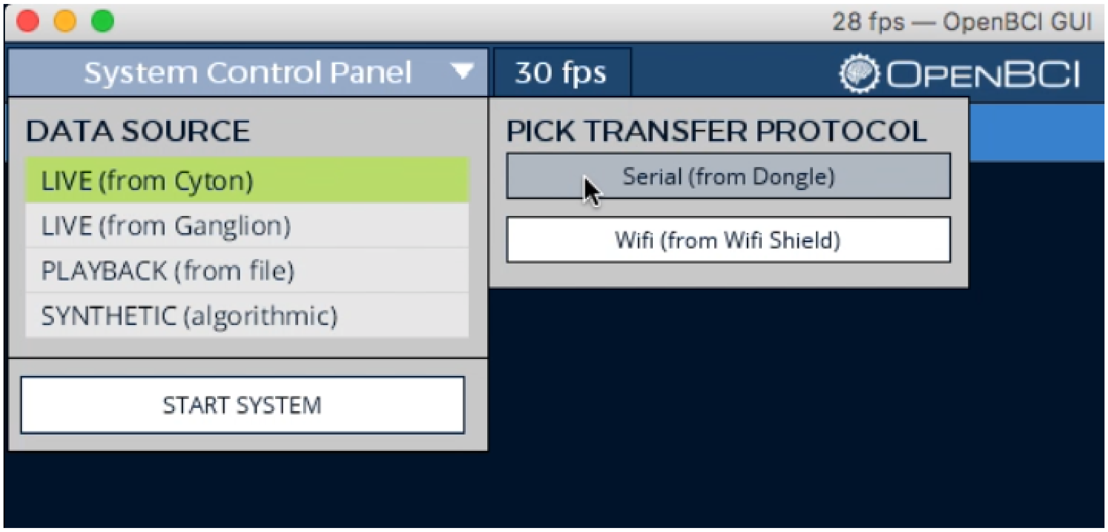


**Figure 2:** Select "Serial (from Dongle)" to retrieve data from the dongle.

3.3 Identify your dongle's Serial/COM port (e.g., /dev/tty for Mac/Linux or COM for Windows) or choose auto-connect (Figure 3).

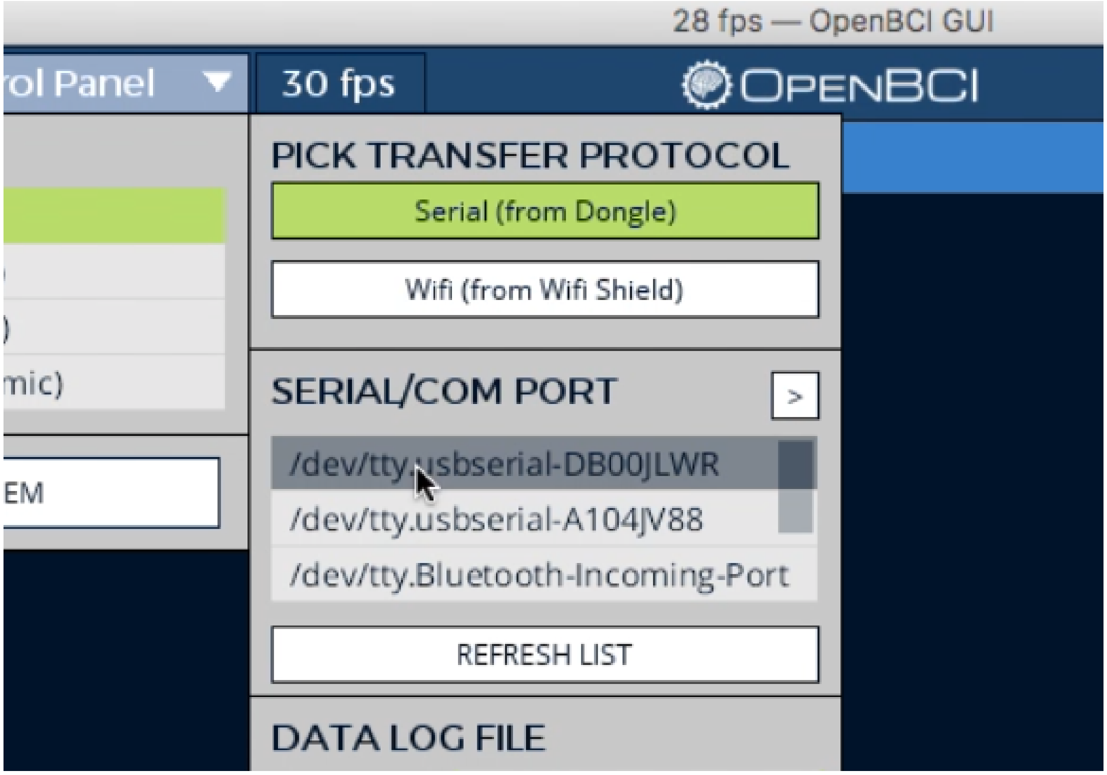


**Figure 3:** Identify your dongle, or in some versions, select "Auto-connect."

3.4 Press the "START SYSTEM" button, and wait ~5 seconds for the GUI to establish a connection (Figure 4).

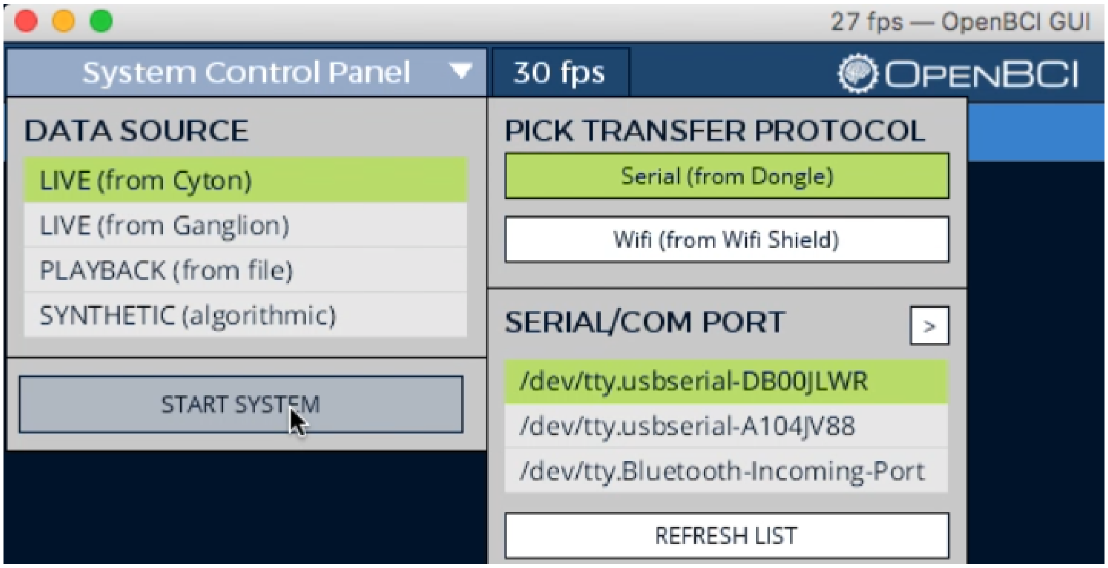


**Figure 4:** Select "START SYSTEM" and wait for the data recording page to load.

3.5 Now that the OpenBCI_GUI is connected to your Cyton, you may press Start Data Stream in the upper left-hand corner (Figure 5).

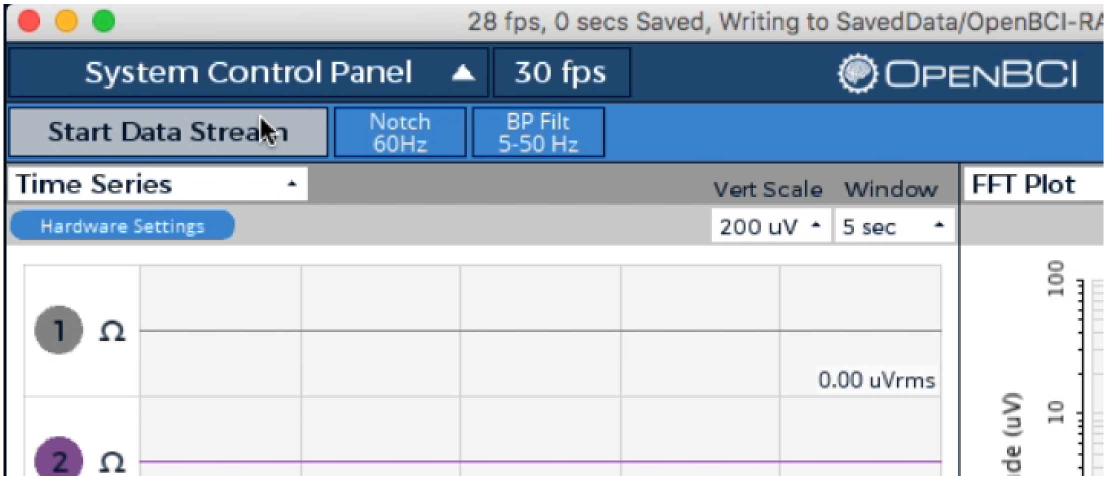


**Figure 5:** Select "Start Data Stream" to begin recording.

3.6 Each connected channel in the Time Series widget should now appear, and the traces on the FFT graph in the upper right will shift upwards instantly (Figure 6).

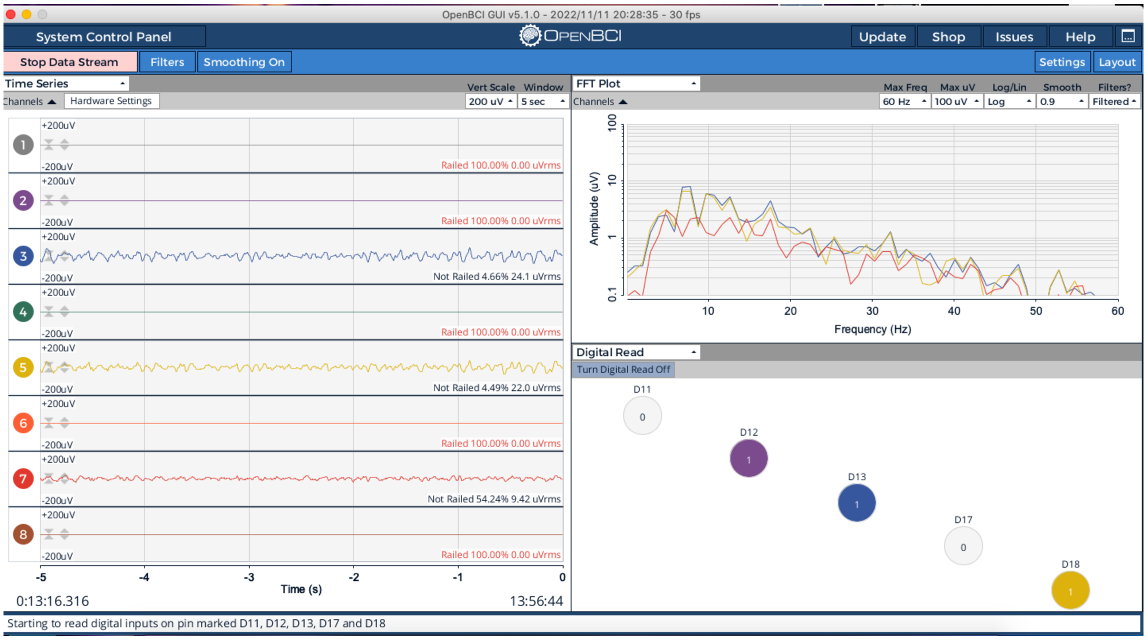


**Figure 6:** From the dropdown menu in the top-left corner, select the required widgets to monitor the signal.

3.7 When you press **Stop Date Stream**, the corresponding files will be automatically saved in both .csv and .txt formats in the OpenBCI GUI directory.

#### Electrode and head post assembely

4 **Electrode preparation:**

- Please cut an approximately 2.5 cm piece of stainless steel wire (Figure 7.1).
- Remove the coat of 0.5 cm of the wire at each end (Figure 7.2).
- Bend the uncoated part of the wire upward and create a hook (Figure 7.3)
- Near the tip of the hook to the body of the electrode (Figure 7.4).
- Use hemostat forceps to lock the electrode's inverted tip close to the body(Figure 7.5).
- Use another hemostat forceps to rotate the loopy part (Figure 7.6).
- After creating a loop, use your finger to bend the electrode into an L-shape (Figure 7.7).

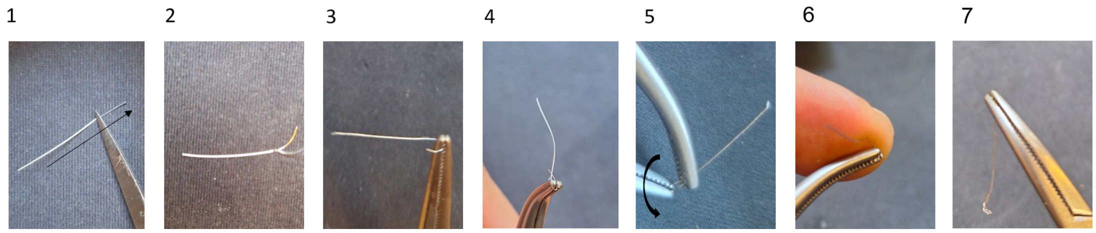


**Figure 7:** Electrod preparation

Preserve the electrodes in a 70% ethanol solution.
Before using the electrodes, ensure that all the alcohol has evaporated. Then, it is recommended that they be washed with normal saline before implantation.
**Consult your veterinarian regarding the regulations and protocols for disinfecting electrodes**¥

5 **Constructing head post:**

To create the head post, use a hypodermic needle and shape it with a hemostat clamp. Blunt, sharp edges of the needle (Figure 8).

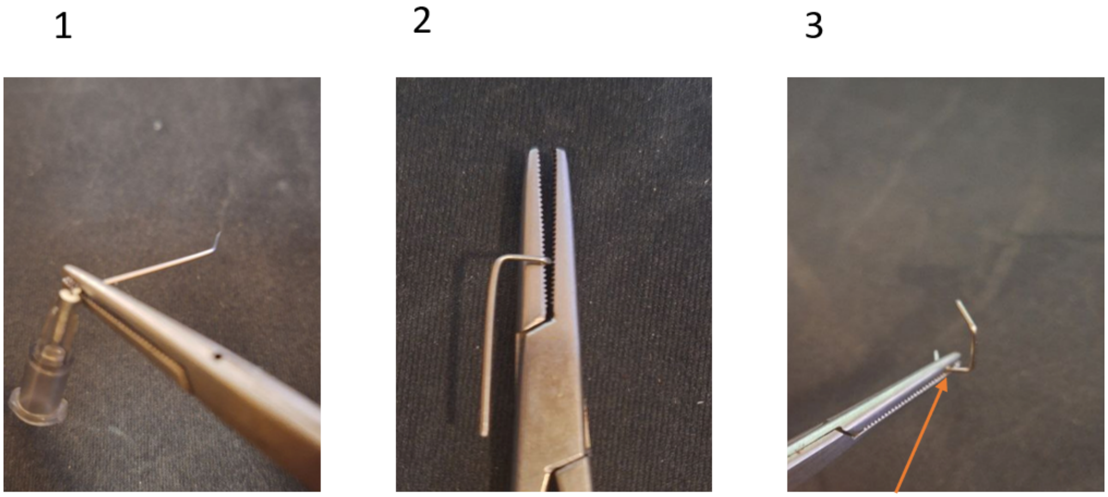


**Figure 8:** Head post preparation steps. Orange arrow shows the base part of the head post.

#### Electrode montage

6 **Electrode montage:**

In both isoflurane levels and auditory presentation experiments, we employed four electrodes: two for reference and noise cancellation and two for signal recordings. In the first experiment, the reference and noise-canceling electrodes were positioned at separate locations on the cerebellum for all finches. In the second experiment, the noise-cancelling electrode was placed in the frontal region, and the reference electrodes were placed in the cerebellum. For the first experiment, we used fixed sites—one in the frontal lobe and another in the auditory cortex.

However, in the second experiment, where precise localization was essential, electrodes were specifically placed in the HVC and CMM regions.

**The electrodes were positioned beneath the skull and above the meninges without any additional conductive materials.** After surgery, all electrodes were connected to the appropriate pins on the OpenBCI board (as detailed in Table 1), and then each experiment was conducted.

**All signal electrodes were referenced to a ground electrode placed on the cerebellum, and no differential signals were compared between pairs of recording electrodes.**

#### Recording iEEG signals

7 **Connecting open BCI board:**

After the surgery is completed and you are ready to start the experiment, position the board and battery close to the anesthetized finch while keeping the Raspberry Pi as far from the isolated box as possible, using a wire to minimize noise. Then, connect all recording electrodes on the finch's head to one of the seven recording pins (N1P to N7P). At this stage, note which electrode is connected to which brain area and try to maintain the same wiring in future experiments. For instance, pin 5 can be connected to the anterior electrode and pin 7 to the posterior electrode. This will facilitate signal analysis. Next, connect the reference and noise-canceling pins to the two remaining electrodes (the two in the cerebellum for experiment one and the anterior and cerebellum for experiment two). Using unoxidized connectors is important for a better signal-to-noise ratio (SNR). To reduce noise, try to shield the finch's surroundings with insulating material.

8 **Impedance check:**

When you launch the OpenBCI GUI and connect to the board as specified in step 3, click on **Cyton Signal** in the dropdown menu at the upper left of the window and check the impedance of the connected electrodes.

The following figure shows the impedance check during the auditory task. HVC (channel 5) displays 12 kOhm, and CMM (channel 7) displays 20 kOhm.

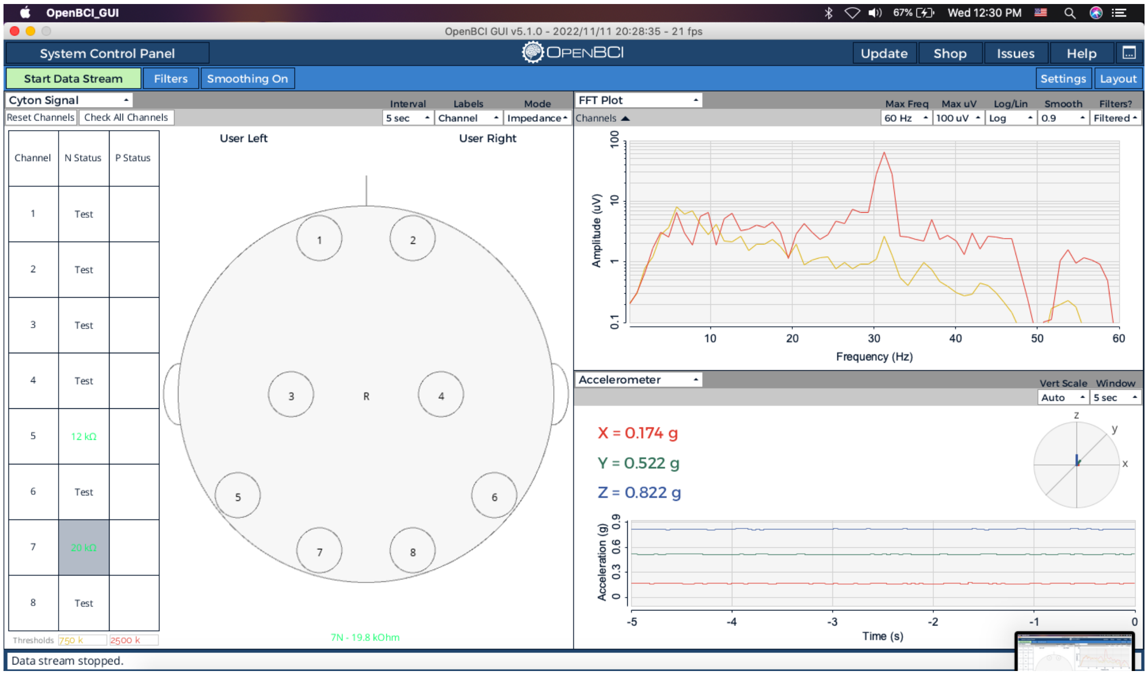


**Figure 9:** Perform an impedance check before starting the auditory experiment.

9 **Anesthetized recording:**

After surgery and electrode implantation, OpenBCI recording was set up, and signals were recorded twice for approximately 120 seconds at each of the following isoflurane concentrations: 2.5%, 2%, 1.5%, 1%, 0.6%, and 0.4%. The data was then extracted based on the varying isoflurane levels for further analysis of suppression duration

10 **Auditory task recording:**

For this part of the experiment, recordings from the HVC and CMM regions were made while five different auditory stimuli were presented: white noise, BOS (bird's own song), conspecific song, reverse BOS, and 5 kHz tones. Spectrograms and event-related potentials for each task condition were analyzed.

#### Stimulus persentation and data synchronization

11 **Stimuli:**

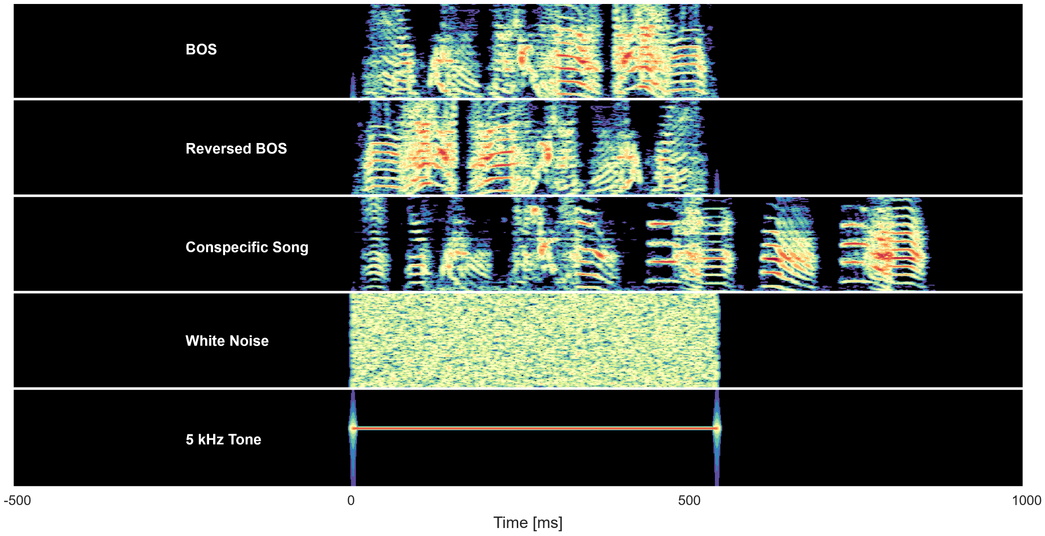


**Figure 10**: Spectrogram of the presented stimuli.

To validate the recorded signals, we presented the subject with five auditory stimuli and collected data from two regions. The stimuli included the bird's own song (BOS), reversed BOS, conspecific song, white noise, and a 5kHz tone (Figure 10), with the BOS recorded 2 days before the experiment. We selected these stimuli because the dorsal brain regions that most influence the recorded signal (HVC and CMM) respond strongest to BOS. Each stimulus was presented 60 times, with intervals randomly chosen between 4 to 6 seconds.

12 **##Shahriar###**

**Stimuli presenting setup:**

""""""""""""""""""""""""""""""""""""""""Write about Raspberry Pi """"""""""""""""""""""""""""""""""""

""""""""""""""""""""""""address the reader to the codes using GitHub repository """"""""""""""""""""""""""""""

13 **Check the Open-BCI Trigger input:**

The input pins on the open BCI board can be used to deliver the external signal. Ensure your Cyton firmware is later than **v3.1.0 by following the tutorial here i**f **the board's firmware is not later than** v3.1.0.

You can check that your board gets the trigger signal by following below steps:

First, open the open-BCI GUI, start data streaming, and open a Digital read window (Figure 11).

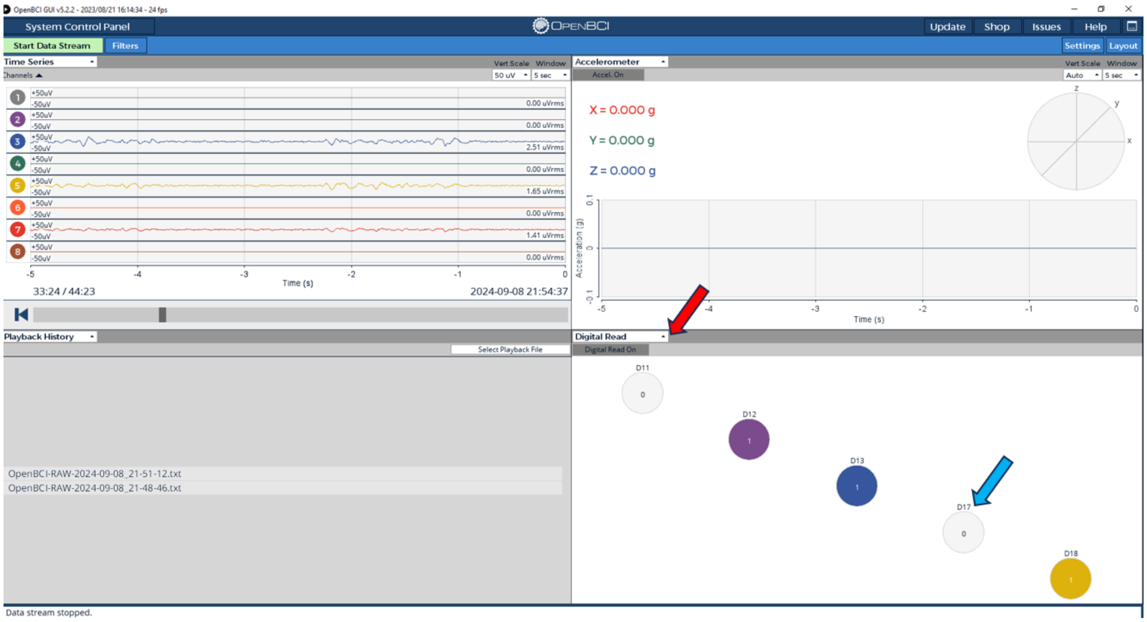


**Figure 11**: Open-BCI GUI: The Red arrow shows the icon you can choose trigger display from, and the blue arrow shows the D17 trigger input position

Then, insert one side of a jumper Wire into the VDD socket on the board. Inset the other head of the wire
to the D17 socket (Figure 12). If the trigger system works, the D17 circle on the GUI will be turned on. Connecting and disconnecting these pins together will turn the D17 circle on and off.

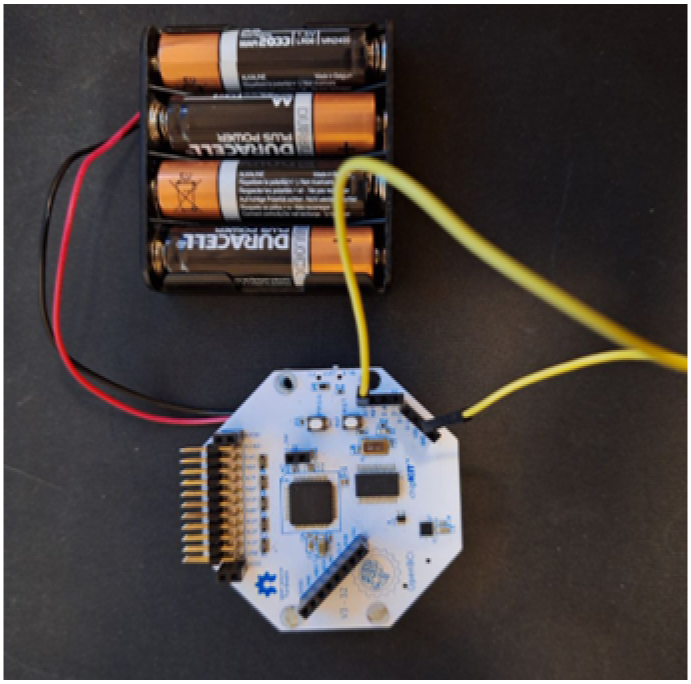

Figure 12: Connecting the VDD to the D17 pins to check the board trigger.

14 **Optocoupler circuit:**

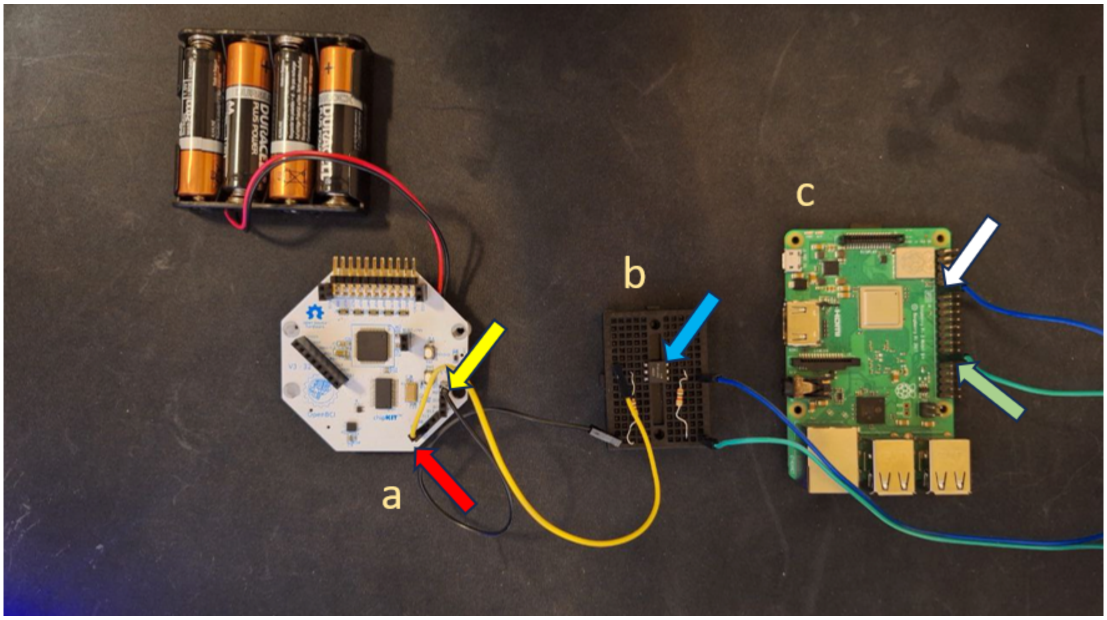


Figure 13: Connection between the Raspberry Pi and open BCI board. a: The Cyton board, VDD pin: red arrow, D17 trigger input: yellow arrow. b: Optocoupler circuit, CNY17F: blue arrow. Raspberry Pi board ground pin: White arrow. Digital write: Green arrow

Connect the Raspberry Pi system to the Open-BCI board using an optocoupler circuit. The Open-BCI site provides a detailed description of the connection between an external trigger and the board here: https://docs.openbci.com/Cyton/CytonExternal/#optoisolation.
We used a very similar optocoupler to what is used on the Open-BCI page, CNY17F, from Vishay https://www.vishay.com/docs/83607/cny17f.pdf. The optocoupler circuit was assembled on a breadboard.
We used a slightly modified version of the circuit. The diagram is provided in **Figure 15.**

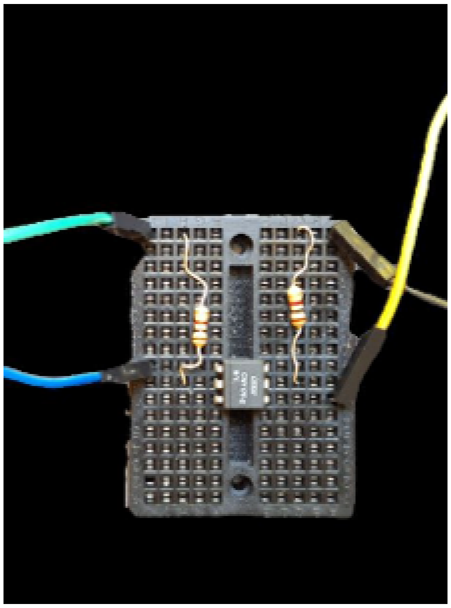

Figure 14: Optocoupler circut

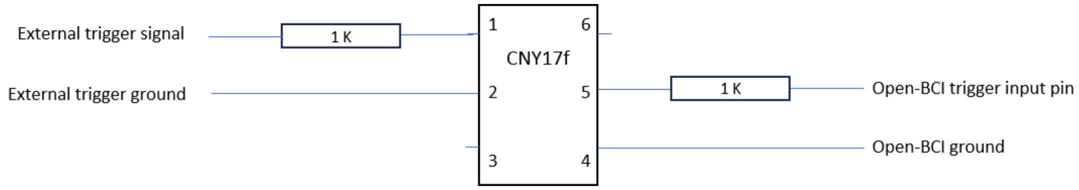


Figure 15: Optocoupler circut diagram

#### Data analysis

15 **##Shahriyar###**

*****You should provide the address for the relevant codes, which Shahriar should do.*********

**Structure of recorded data and analysis method:**

The data was processed in Python, primarily utilizing the MNE package. First, the raw data files, formatted as .txt, were loaded, and the relevant columns representing EEG signals from two channels, along with a trigger channel for marking stimuli, were extracted and organized for further analysis.

The data was filtered using a bandpass filter between 1 and 40 Hz to isolate the frequencies of interest. After filtering, the timing of events related to the auditory stimuli was detected, and these events were used to classify the different stimulus types. Each stimulus event was detected using a 5-sample time window to ensure accurate data segmentation.

Once the events were identified, epochs were created for each stimulus presentation, with a time window starting one second before stimulus onset and spanning three seconds after. A baseline correction was applied to remove any drift or noise from the data before stimulus onset.

The epochs were then averaged over time to represent the Event-Related Potential (ERP) corresponding to each of the five auditory stimuli and computed the Time-Frequency Representation (TFR) using the Stockwell transform method (Stockwell 2007; Moukadem et al. 2014).

16 **##Shahriyar###**

*****You should provide the address for the relevant codes, which Shahriar should do.*********

We utilized a real-time algorithm developed by Westover et al. in 2013 that automatically distinguishes between suppressions and bursts in EEG readings (Westover et al., 2013). This enabled us to rely on the total suppression duration for each task condition and conduct statistical comparisons of these durations. For each level of isoflurane (low, medium, and high), where low is defined as below 1%, medium as below 2%, and high as 2% and above (Figure 16), we calculated the suppression duration for ten blocks of 120-second recording segments and performed statistical analyses to compare the conditions.

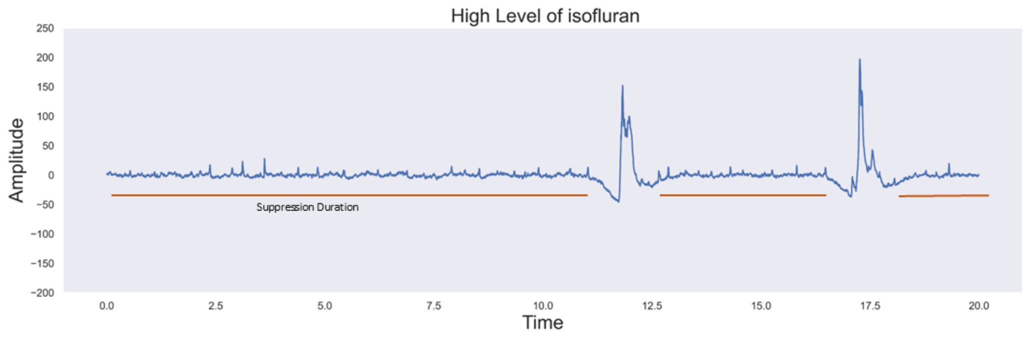


**Figure 16:** Example of suppression duration in high level of isoflurane (crop of a signal for 20 s)

17 **Auditory evoked activities during anesthesia:**

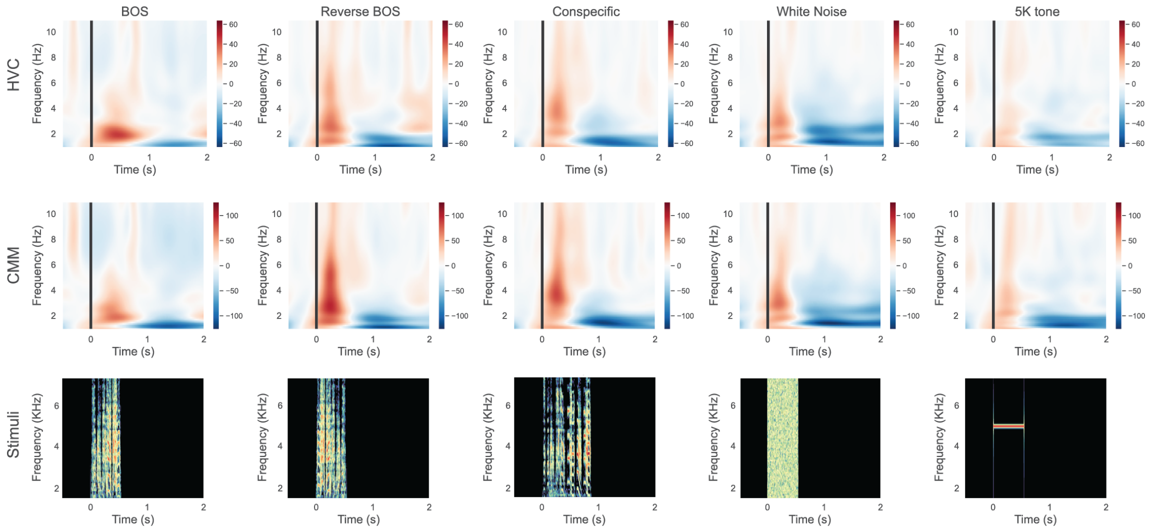


***** Caption is needed****

The recorded signal was first bandpass filtered from 1 to 30 Hz, and then each channel was z-scored using a baseline period. The signal for each trial was transformed to the frequency domain using continuous 1-D wavelet transform, and trials were averaged by stimulus type. To reduce noise, we averaged the frequency domain signal using a 3-second sliding window with 30% overlap, then subtracted this from each stimulus response.
The plot above shows a neural response in the delta band in both channels, occurring only in response to the BOS, which aligns with the literature indicating that both dorsal nuclei, as higher-level regions, predominantly respond to BOS.

#### Bird Handling and Surgical Process:

18 **Anesthesia mask[¥]:**
We designed and tested a 3D printable adaptor for gas delivery and head fixation under the supervision of a veterinarian. It consists of two parts. Please check the regulations in your

institute regarding the use of the anesthesia method and seek consultation from your veterinarian.

PLA printed materials cost around 20 $

funnel.stl 1.2MB base (1).stl 828KB

19 **Constructing ear bar adaptor¥:**

Note: There are better solutions for specific stereotaxic device adaptors for birds on the market. You can find one example here **https://kopfinstruments.com/product/model-914-small-bird-adaptor-and-model-914t-small-bird-surgical-head-holder/** . Please check the available options and seek consult from your veterinarian. We developed our method for creating air bars since we couldn't access the usual solutions. Our veterinarian approved our method. **Two of the animals were fixed with suitable earbars without adaptors.**

1 2 3 4 5

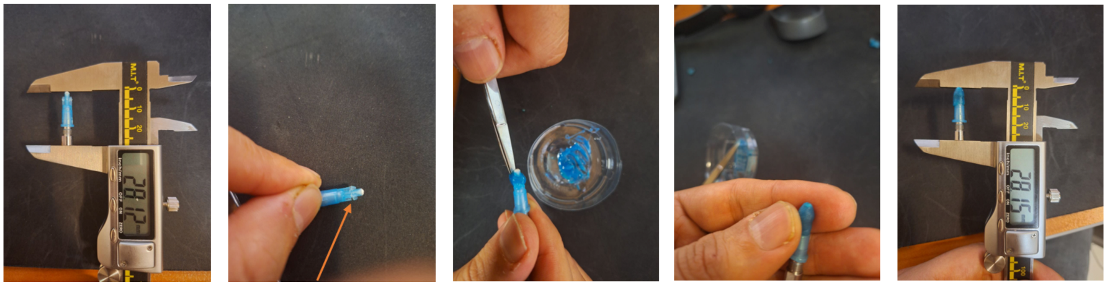

Preparing ear-bar adoptor. Orange arrow shows the postion of notches

- Detach the hypodermic needle from its plastic base.
- Blunt the tip of the detached part to remove its sharpness and inspect the tip to ensure it's not sharp.
- Measure the length of both adapters and ensure their sizes are equal. (Figure ... 1)
- As shown in the figure, create notches along the base's length and measure its length with a clipper. Both adapters should be the same size. (Figure ... 2).
- Prepare the silicone mixture and wrap it around the tip of the needle base, ensuring that the silicone paste fills the notches (Figure ... 3) . We used condensation silicone from Coltene Inc.; you can find additional information here: https://www.basiqdental.nl/en_NL/p/103374.
- Shape the silicone with your finger to form a pointy tip. (Figure ...4)
- Measure the length of both adapters and ensure their sizes are equal. (Figure ... 5)

20 **Animal ¥:**

Female or male zebra finches older than 60 days are suitable for this operation.

**Do not use underweighted animals. They have a lower survival rate under anesthesia.**

21 **Preoperative procedures ¥:**

Restrict food intake before the operation andwater consumption before the surgery.

**Do not restrict feeding to more than 30 minutes.**

22 **Anesthesia induction ¥:**

Place the animal inside the induction chamber. Start with 2 minutes of oxygen delivery and then increase the gas concentration by 0.5 percent every 1 minute until it reaches 2 percent. Keep the animal under the gas chamber for two minutes. Administer Ringer Lactate every 20 minutes throughout the operation.

**Safety information**

Isoflurane leakage from anesthesia machines can affect experimenters' health. To avoid this issue, use a suitable scavenging system. Staff should be well informed and educated about the possible hazards of anesthetic gas.

23 **Preparation of the incision site and skull for craniotomy:**

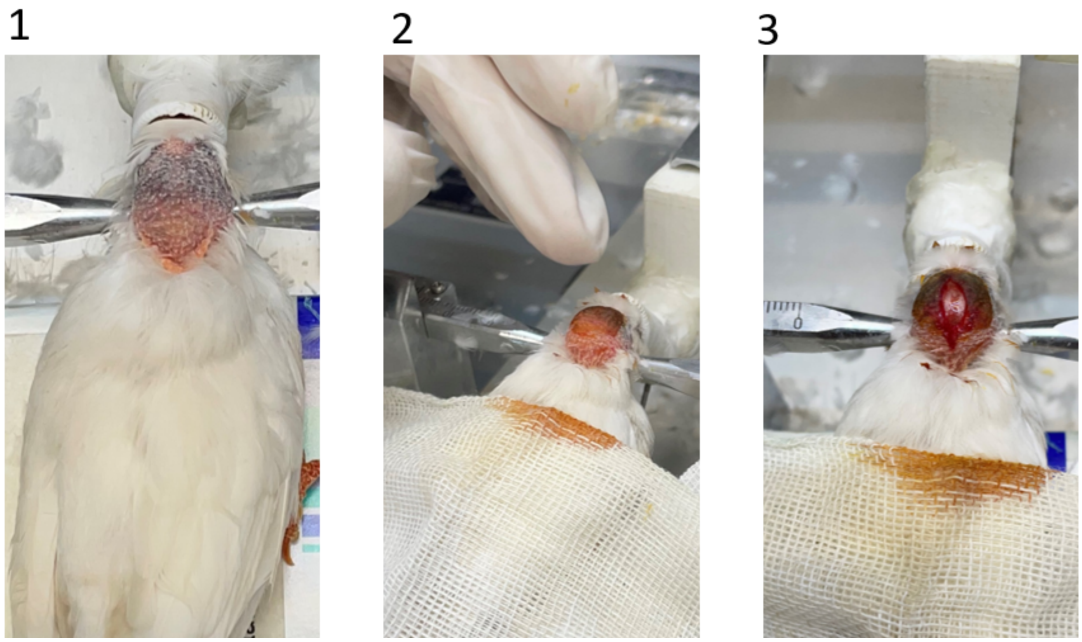


Preparation of scalp and create an incision on it.

- Gently remove the covering feathers from the skull.
- Use the scalp as an incision and subcutaneous injection site. Disinfect the surface of the skull by applying diluted betadine and injecting lidocaine subcutaneously. Wait for the drug to be absorbed.
- Make a midline incision between the eyes and the cerebellum. Retract the edges of the incision. The bifurcation site should be immediately recognizable.
- Scrape the skull to remove the connective tissue using a tiny piece of sterilized tampon.
- Allow the skull to dry.

24 **Localizing the electrode sites:**
After making the incision.
- Locate the bifurcation and gently scrape the outer layer of the skull bone and cerebellum at the bifurcation site.
- Hydrate the area to see beneath the inner layer of bone.

- Drill two holes in the outer layer of the bone on the cerebellum, one millimeter caudal and one millimeter lateral to the bifurcation area.
- You can attach a needle to the stereotaxic, scratch the skull to mark the recording sites and drill a hole.

Coordination of recording sites:

ANT: AP: 5 mm, ML: 2 mm

"""""""""""""""""""" **Amirreza** """""""""""""""""""""""""""""""""""

HVC: 0.2 mm AP | 2.3 mm ML

CMM: 2 mm AP | 0.4 mm ML

25 **Craniotomy¥:**

We used a 1 mm ball-shaped dental burrel with a surgical micromotor to drill the skull. Electrodes were placed underneath the skull on the brain's surface on dura matter.

**Drilling sites must be continuously hydrated with normal saline to prevent neural damage from frictional heating.**

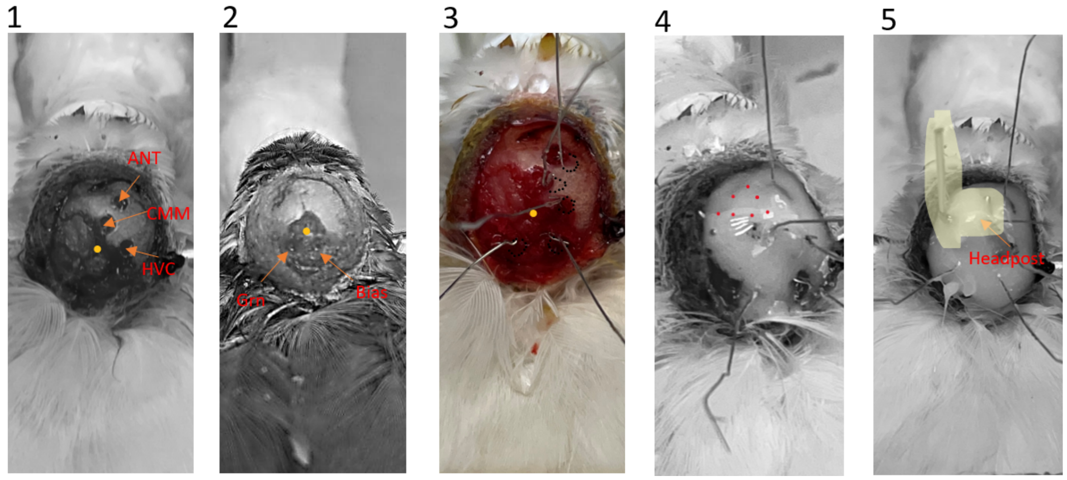


Craniotomy and electrode replacement: Three recording electrodes are placed on CMM (caudomedial mesopallium) and HVC. The ANT (anterior) area is not put in the auditory area.

The yellow dot represents the bifurcation location. In Figure 2, the location of the Ground and Bias electrodes is shown. In Figure 3, black dotted circles show the placement of the electrode contacts. Red dots indicate the location of the head post base in Figure 4. The base of the head post is placed perpendicular to the skull inside the forming cement. The orange arrow indicates the base of the headpost, and the highlighted area indicates the headpost.

26 **Electrode placement and fixation:**

All of the electrodes are placed under the skull on the dura mater. First, insert the loop part into the skull hole (Figure 1), then push the electrode into the hole's edge, ensuring that the electrode body is connected to the skull (Figure 2). A cartoon picture illustrating the position of the electrodes related to the skull and brain is provided. After inserting the electrodes into place, we locally administered antibiotics using Ciprofloxacin eye drops (1 drop of 0.3% solution)¥. We then waited for the water on the surface of the exposed area to evaporate. Next, we applied glass ionomer cement to the skull and the drilled holes. The cement filled the cavities, and once it solidified, the electrodes were securely fixed in position.

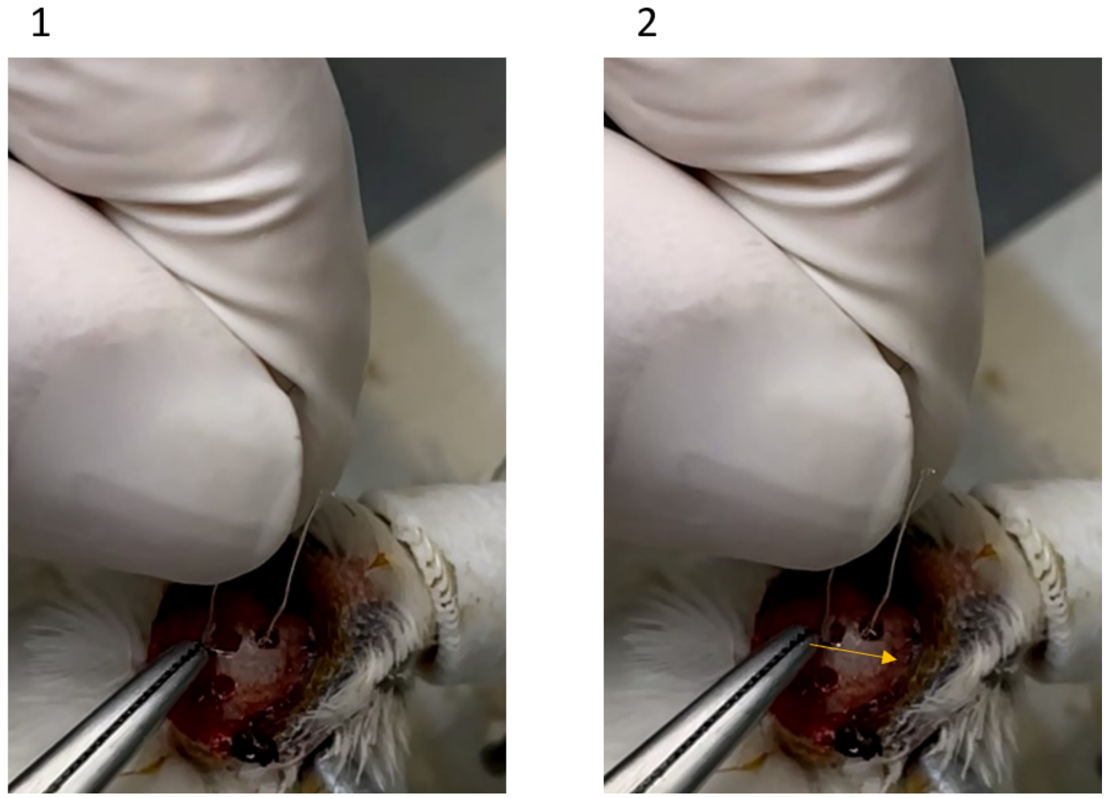


Elctrode placemetn in recording sites. the yellow arrow shows the direction of the force that is applied on the electrode body to be placed under the skull

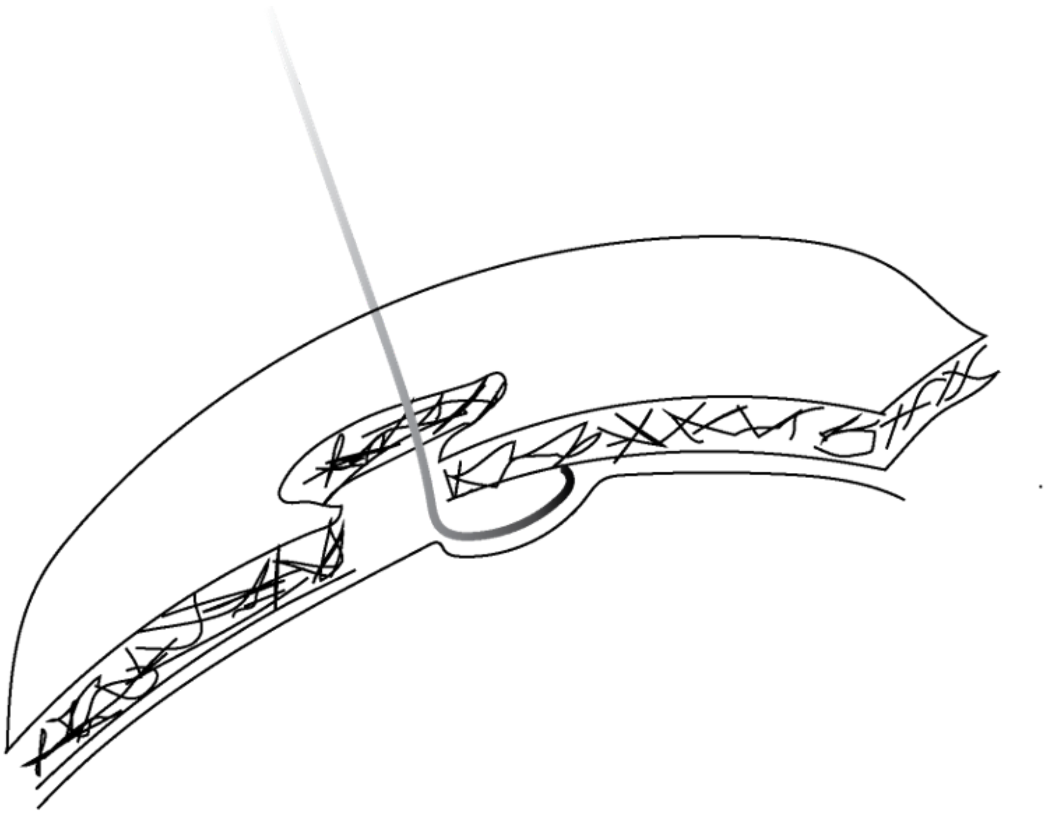

All electrodes are placed under the skull and above the meninges. The force between the skull and brain fixes the electrode automatically if pushed into this position.

27 **Head Post implantation:**

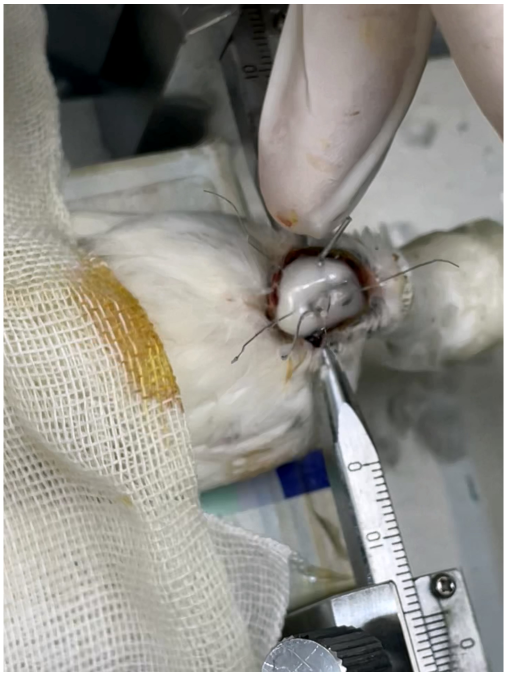

Implanting headpost

28 **Post-operative care¥ :**

Make sure that the cement is completely cured. After that, the animal is transferred to its private cage and used as a heat source to keep its body warm. We use a ventilator heater to maintain the temperature during recovery and surgery.

**Note that ventilator heaters can increase the rate of water loss through evaporation.**

29 **Adaptation period¥ :**

Monitor the bird during the 7-day adaptation period to detect any adverse effects. Usually, birds adapt well to implanted materials during this period. Some adverse neurological problems, like movement disorders and hearing or vision loss, may occur. Check the animals to exclude them if these symptoms are present. Ask your veterinarian for help diagnosing these possible problems.

30 **Animal housing¥ :**

Avoid placing birds in multiple cages; they try to cut their friends' electrodes. Place their cages close to each other to avoid the stress of social isolation.

#### Troubleshooting

31 **Problem:** The trigger system is not working when you check it following step 49

**Solution:** Update the Cyton board firmware. All related information is available here: **https://docs.openbci.com/Cyton/CytonProgram/**.

**Problem**: The Optocoupler circuit is not working or causing noise in the signal:

**Solution:** Check the connections. Use a soldered circuit instead of a breadboard for a more reliable connection.

**Trouble**: The uncoated end of the electrodes was accidentally covered by dental cement.

**Solution**: Wait for the cement to be cured, then use a hemostat clamp to smash the covered part.

**Trouble**: The electrode gets accidentally contaminated by contact with a non-sterile surface.

**Solution**: repeat the disinfection process and re-implant the electrode.

**Trouble**: The electrode gets accidentally contaminated by contact with a non-sterile surface.

**Solution**: repeat the disinfection process and re-implant the electrode.

**Trouble**: The not-railed of the time series recording.

**Solution**: When the signal is noisy or saturated, it may increase and sometimes fail to record at 100%, not a railed signal. If this occurs, reconnect to the board, replace the connectors for the affected channel, and check both the reference and noise-canceling pins.

**Trouble:** Insufficient Power Supply During Recording

**Solution**: Before recording, ensure your AA batteries are fully charged to prevent interruptions. Check battery levels and carry spares for longer sessions.

#### Protocol references

1. Brandon Westover M, Shafi MM, Ching S, Chemali JJ, Purdon PL, Cash SS, Brown EN. Real-time segmentation of burst suppression patterns in critical care EEG monitoring. J Neurosci Methods. 2013 Sep 30;219(1):131-41. doi: 10.1016/j.jneumeth.2013.07.003. Epub 2013 Jul 23. PMID: 23891828; PMCID: PMC3939433.

2. Stockwell RG. Why use the S-transform? In: Rodino L, Schulze BW, Wong MW, eds. Pseudo-Differential Operators: Partial Differential Equations and Time-Frequency Analysis. Vol 52. Fields Institute Communications. Providence, RI: American Mathematical Society; 2007:279–309. doi:10.1090/fic/052.

3. Moukadem A, Bouguila Z, Ould Abdeslam D, Dieterlen A. Stockwell transform optimization applied on the detection of split in heart sounds. In: Proceedings of EUSIPCO-2014. Lisbon; 2014:2015–2019. IEEE. Available at: https://ieeexplore.ieee.org/document/6952743.
